# Spontaneous selection of *Cryptosporidium* drug resistance in a calf model of infection

**DOI:** 10.1101/2021.01.04.425361

**Authors:** Muhammad M. Hasan, Erin E. Stebbins, Robert K.M. Choy, J. Robert Gillespie, Eugenio L. de Hostos, Peter Miller, Aisha Mushtaq, Ranae M. Ranade, José E. Teixeira, Christophe L. M. J. Verlinde, Adam Sateriale, Zhongsheng Zhang, Damon M. Osbourn, David W. Griggs, Erkang Fan, Frederick S. Buckner, Christopher D. Huston

**Affiliations:** Cellular, Molecular and Biomedical Sciences Graduate Program, University of Vermont, Burlington, VT, USA; Department of Medicine, University of Vermont Larner College of Medicine, Burlington, VT, USA; PATH, San Francisco, CA, USA; Department of Medicine, University of Washington, Seattle, WA, USA; Department of Biochemistry, University of Washington, Seattle, WA, USA; The Francis Crick Institute, London, UK; Department of Molecular Microbiology and Immunology, Saint Louis University, St. Louis, MO, USA

## Abstract

The intestinal protozoan *Cryptosporidium* is a leading cause of diarrheal disease and mortality in young children. There is currently no fully effective treatment for cryptosporidiosis, which has stimulated interest in anticryptosporidial development over the last ∼10 years with numerous lead compounds identified including several tRNA synthetase inhibitors. In this study, we report the results of a dairy calf efficacy trial of the methionyl-tRNA (*Cp*MetRS) synthetase inhibitor **2093** and the spontaneous emergence of drug resistance. Dairy calves experimentally infected with *Cryptosporidium parvum* initially improved with **2093** treatment, but parasite shedding resumed in two of three calves on treatment day five. Parasites shed by each recrudescent calf had different amino acid altering *CpMetRS* mutations, coding either an aspartate 243 to glutamate (D243E) or a threonine 246 to isoleucine (T246I) mutation. Transgenic parasites engineered to have either the D243E or T246I *Cp*MetRS mutation using CRISPR/Cas9 grew normally but were highly **2093** resistant; the D243E and T246I mutant expressing parasites respectively had **2093** EC_50S_ of 613- or 128-fold that of transgenic parasites with wild-type *Cp*MetRS. In studies using recombinant enzymes, the D243E and T246I mutations shifted the **2093** IC_50_ by >170-fold. Structural modeling of *Cp*MetRS based on an inhibitor-bound *Trypanosoma brucei* MetRS crystal structure suggested that the resistance mutations reposition nearby hydrophobic residues, interfering with compound binding while minimally impacting substrate binding. This is the first report of naturally emerging *Cryptosporidium* drug resistance, highlighting the need to address the potential for anticryptosporidial resistance and establish strategies to limit its occurrence.

**Importance:** *Cryptosporidium* is a leading protozoan cause of diarrhea in young children with no reliable treatment. We report results of a dairy calf drug efficacy trial and the spontaneous emergence of drug resistance. *Cryptosporidium parvum* infected calves initially improved with drug treatment, but infection relapsed in two animals. Parasites shed by each recrudescent calf had mutations in the gene encoding the drug target that altered its amino acid sequence. Recapitulation of the drug target mutations by CRISPR/Cas9 genome editing resulted in highly drug-resistant parasites, and recombinant mutant enzymes were resistant to inhibition. This is the first report of naturally emerging *Cryptosporidium* drug resistance. There is a currently a great opportunity to impact public health with new drugs to treat cryptosporidiosis, and this report highlights the need to address the potential for anticryptosporidial resistance and establish strategies to limit its occurrence in order to realize their full potential.

**One-sentence summary:** Drug-target point mutations mediating anticryptosporidial resistance spontaneously arose in the dairy calf *C. parvum* infection model.

## Introduction

Infection of small intestinal epithelial cells with *Cryptosporidium* parasites causes the diarrheal illness cryptosporidiosis that in humans is mostly due either to *Cryptosporidium hominis* or *Cryptosporidium parvum* (*1-4*). Symptoms of cryptosporidiosis last twelve days on average in immunocompetent people but are often prolonged and potentially fatal in malnourished children and immunocompromised people, such as those with AIDS and transplant recipients (*1*). In addition to being a prominent cause of chronic diarrhea in AIDS patients, cryptosporidiosis was a leading cause of life-threatening diarrhea in children under two years of age in a landmark case-control study recently conducted at seven high-burden sites in Africa and the Indian subcontinent (*5, 6*). *Cryptosporidium* infections are also strongly associated with childhood malnutrition, growth stunting, and delayed cognitive development (*7, 8*).

Despite the large public health burden, no *Cryptosporidium* vaccine exists and treatments for cryptosporidiosis remain inadequate. Nitazoxanide is the only FDA-approved drug for cryptosporidiosis and is reliably efficacious only for immunocompetent adults for whom it shortens the duration of diarrhea by 1-2 days (*9*). Unfortunately, nitazoxanide is no more effective than a placebo in HIV positive people, and data for young malnourished children are limited to small studies in which it appears at best modestly efficacious (*10, 11*).

The need for improved anticryptosporidials motivated multiple drug development efforts employing both phenotypic and target-based methods over the last decade. These efforts have led to the identification of numerous promising lead compounds and two pre-clinical candidates that are poised for human studies (i.e. a Novartis phosphatidyl-inositol-4-kinase inhibitor and the 6-carboxamide benzoxaborole AN7973)(*12-20*). There is the potential to realize an enormous public health benefit with intelligent deployment of improved anticryptosporidial drugs.

In many ways, the anticryptosporidial field evokes the situation for malaria shortly before the introduction of chloroquine in 1945, a time at which there was nearly a blank slate for treating malaria and the inevitability of drug resistance was not appreciated. Yet, chloroquine resistance first became evident for malaria around 1957, just twelve years after its release (*21*). Based on the hard lessons from malaria chloroquine resistance and antimicrobial resistance in general, combination therapy is now the standard for antimalarial development, as well as for treatment of numerous other infectious diseases where the risk of resistance is high (e.g. HIV and tuberculosis) (*22*). This history stresses the need to understand the potential for *Cryptosporidium* drug resistance and how to best deploy new anticryptosporidials. To date, the possibility of anticryptosporidial resistance remains completely unexamined.

Aminoacyl-tRNA synthetase (aaRS) inhibitors are protein synthesis inhibitors with promise as potential treatments for bacterial, fungal, and parasitic infections. AaRS enzymes catalyze the formation of tRNAs charged with their corresponding amino acids that serve as the substrates for protein synthesis. Two aaRS inhibitors are already approved for human use: the topical anti-staphylococcal antibiotic mupirocin, a natural product that inhibits isoleucyl-tRNA synthetase (*23*); and tavaborole, a leucyl-tRNA synthetase inhibitor that is approved for topical treatment of onychomycosis (*24, 25*). Beyond these topical agents, several aaRS inhibitors are in clinical trials as systemic antimicrobials, including a methionyl-tRNA synthetase (MetRS) inhibitor for treatment of *Clostridium difficile* infections (*26*), a leucyl-tRNA synthetase (LeuRS) inhibitor for treatment of Gram-negative bacterial infections (*27, 28*), and a LeuRS inhibitor for treatment of multidrug-resistant tuberculosis (*29, 30*).

AaRS inhibitors figure prominently within the anticryptosporidial developmental pipeline (*31*). Halofuginone, a prolyl-tRNA synthetase inhibitor, is approved for prevention and treatment of *C. parvum* infection in dairy cattle in Europe, but cannot be used in humans due to a narrow safety window (*32*); other prolyl-tRNA synthetase inhibitors are under study (*16*). Phenylalanyl-tRNA synthetase (PheRS) and Lysyl-tRNA synthetase (LysRS) inhibitors with promising activity in several mouse models of infection are also under development (*14, 33*). We previously reported that a MetRS inhibitor, **2093**, potently inhibits the *Cryptosporidium* MetRS enzyme and is efficacious against *C. parvum* with no signs of toxicity in two murine models of infection (*34*).

MetRSs are categorized into two forms based on sequence similarity: type I MetRSs (MetRS1) that are homologous to the human mitochondrial MetRS, MetRSs of most Gram-positive bacteria, and those of some parasites, including *Trypanosoma* species; and type II MetRSs (MetRS2) that are homologous to the human cytoplasmic MetRS and those found in most Gram-negative bacteria (*35*). Some Gram-positive bacteria, such as *Bacillus anthracis* and some *Streptococcus pneumoniae* strains, have both a MetRS1 and a MetRS2. *C. parvum* and *C. hominis* each have a single *Cryptosporidium MetRS* gene that encodes a protein homologous to MetRS1 enzymes. We identified the anticryptosporidial activity of **2093** and related compounds by screening MetRS1 inhibitors under development for treatment of Gram-positive bacterial infections and trypanosomes (*34*).

We now report results of a **2093** drug efficacy trial performed in *C. parvum* infected dairy calves and the spontaneous emergence of MetRS inhibitor resistance. We identify two mutant parasite strains with different single amino acid substitutions in the *C. parvum* MetRS. We confirm *C. parvum* resistance to **2093** and further investigate the mechanism of resistance using a combination of CRISPR/Cas9 genome editing to modify the *C. parvum MetRS* (*CpMetRS*), in vitro enzymatic assays performed with recombinant mutant enzymes, and structural modeling. This study is the first report of emergent *Cryptosporidium* drug resistance and, therefore, highlights a dire need to anticipate the potential for *Cryptosporidium* drug resistance and establish strategies to mitigate its development.

## Results

### Relapse of *C. parvum* infection during MetRS inhibitor treatment of infected dairy calves

Neonatal dairy calves are highly susceptible to *C. parvum* infection and develop a self-resolving diarrheal illness that closely resembles cryptosporidiosis in young children. Experimentally infected calves therefore provide a natural clinical model of *C. parvum* infection to determine both the microbiologic and clinical efficacy of drug leads (*36, 37*). Based on promising results in both *ifn-γ* ^*-/-*^ mice and NOD SCID gamma mice (*34*), we selected the MetRS inhibitor **2093** for testing in the calf model. We previously showed in a cell-based assay that the rate of *C. parvum* elimination by **2093** is maximized by exposure to concentrations at or above three times the EC_90_ (*34*). A preliminary pharmacokinetics (PK) study performed on uninfected calves demonstrated plasma and fecal levels in excess of three times the EC_90_ for over 24 h following a single 10 mg per kg oral dose (Supplemental Fig. 1a). To ensure **2093** levels in excess of three times EC_90_ despite possible reduced absorption and accelerated intestinal transit due to diarrhea, a dose of 15 mg per kg every 12 h was chosen for the efficacy study.

Bull Holstein calves were infected with ∼5×10^7^ *C. parvum* (Bunch Grass Farm Iowa isolate) at 24-48 h of age. **2093** or vehicle control treatments were begun on day two of infection, the time typically corresponding to the onset of diarrhea in this model (*19, 37*), and continued for seven days (Fig. 1a). Figure 1b shows the rate of oocyst shedding in the feces vs. time for individual calves. **2093** initially appeared effective with progressive elimination of oocyst shedding from all treated calves during the first four days of infection. Total oocyst shedding, calculated as the area under the curve for each animal, was reduced nearly two logs during the first four days of compound treatment (Fig. 1c). However, two of three **2093**-treated calves relapsed with a progressive rise in fecal oocysts beginning on day four of treatment; the remaining calf appeared to be cured (Fig. 1b). Recrudescence could not be attributed to inadequate dosing or altered drug levels in the presence of *Cryptosporidium* infection and diarrhea, since both fecal and plasma levels were well in excess of three times the EC_90_ beginning within 5 h of initiating treatment (Supplemental Fig. 1b). These data suggested potential emergence of **2093**-resistant parasites. It was impossible to isolate viable oocysts and test directly for drug resistance in vitro, because recrudescent *C. parvum* shedding was not recognized by qPCR until after sample freezing and euthanizing the animals.

**Fig 1.**
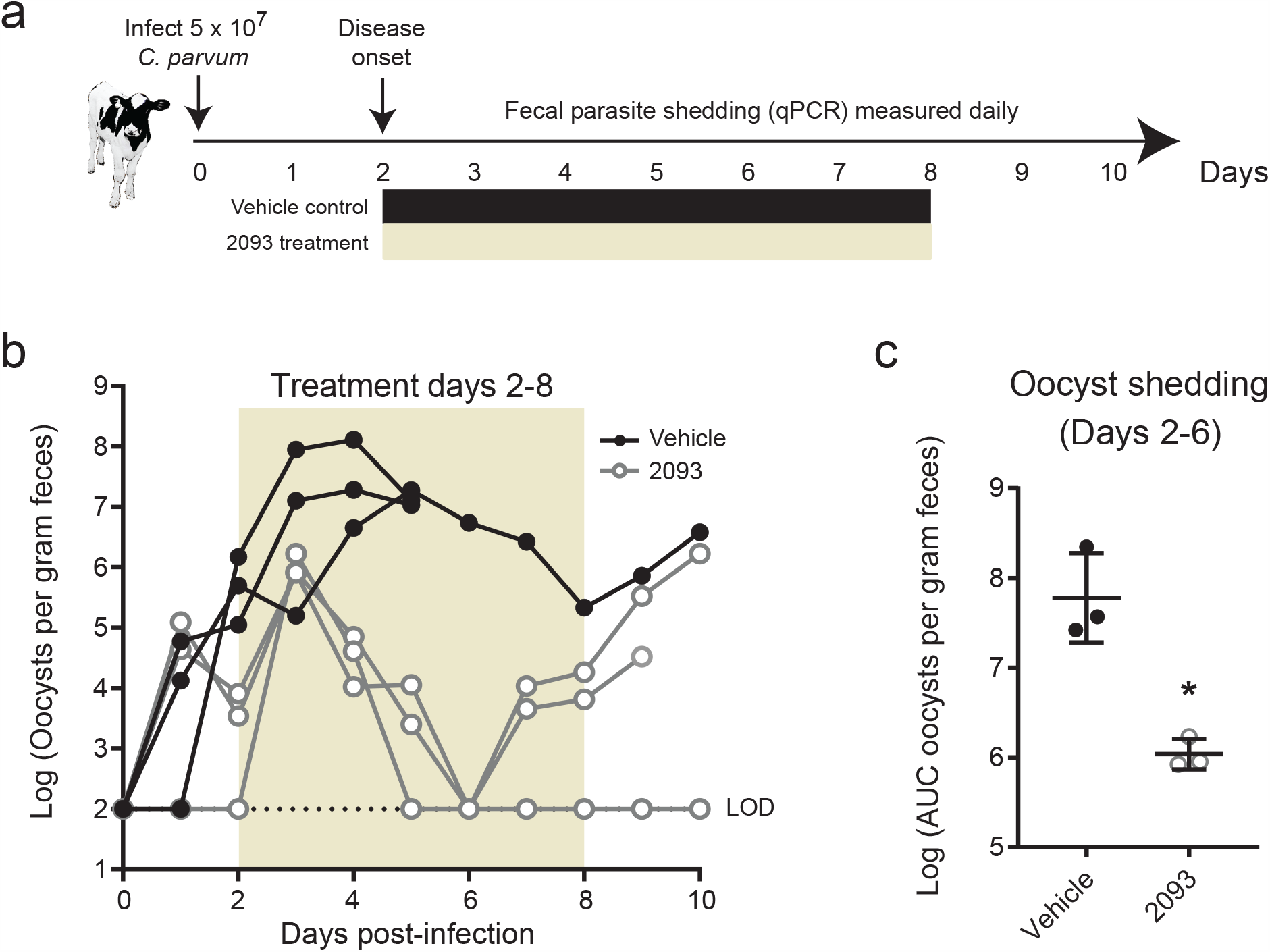
Resurgent *C. parvum* shedding occurred during compound 2093 treatment of infected dairy calves. **(a)** Summary of neonatal calf model and compound **2093** clinical efficacy study. Bull Holstein calves were infected within 48 h of birth by oral administration of ∼5 × 10^7^ *C. parvum* oocysts (Bunch Grass Farms Iowa isolate). Fecal oocyst shedding was quantified daily using qPCR. Vehicle or **2093** (15 mg per kg every 12 h) were administered orally on study days 2-8. **(b)** Oocyst shedding vs. time for **2093**-treated and control calves. The colored box highlights the timing of treatment, and oocyst shedding is plotted for individual animals (n=3 for each group). Note that 2 animals in the control group died on day 5 due to unrelated causes (one due to congenital bladder obstruction and one due to pneumonia). LOD indicates the limit of detection for the qPCR assay. Samples with negative PCR results are plotted at the LOD. **(c)** Total oocyst shedding per gram of feces during study days 2-6. The area under the curve (AUC) in excess of the qPCR LOD is plotted for each calf. Lines show mean and standard deviation. * indicates p = 0.005 (unpaired two-tailed t-test) for reduced oocyst shedding in **2093**-treated animals up to the point of recrudescence in two of the three treated calves.

### *MetRS* gene coding mutations in shed oocysts from 2093-treated calves

Mutation of the target gene is one potential mechanism of acquired drug resistance, so the known target of **2093** provided an opportunity to investigate this possibility. For this, we used total DNA isolated from the feces of each vehicle and **2093**-treated calf on experiment days 1 and 9 as PCR template, and Sanger sequenced the *CpMetRS* genomic locus. Predictably, no PCR product could be obtained from the cured calf on day 9. The genomic *CpMetRS* sequences on day 1 of the experiment were identical to the *C. parvum* Iowa strain gene sequence for all calves, but on day 9 each of the two relapsed calves had a distinct coding single nucleotide variant (SNV) in the *CpMetRS* gene (Supplemental Fig. 2). These variants were not present in the two control calves on day 9. The identified *CpMetRS* mutations were predicted to result in substitutions at amino acids 243 and 246 of the *Cp*MetRS enzyme: aspartate 243 mutated to glutamate (D243E), and threonine 246 mutated to isoleucine (T246I) (Supplemental Fig. 2a). **2093** treatment in the calf model therefore appeared to select for *Cryptosporidium* strains with two independent *CpMetRS* mutations that potentially conferred drug resistance.

### *CpMetRS* coding mutations confer parasite resistance to inhibition by 2093

We used a recently described method for CRISPR/Cas9 genome editing in *C. parvum* (*38*) to determine the effects of the amino acid altering *CpMetRS* mutations that were identified in the calf drug efficacy trial on parasite sensitivity to **2093**. This method combines in vitro electroporation of a plasmid encoding the guide RNA and Cas9 enzyme, and a linear repair cassette, followed by infection of *ifn-γ* ^*-/-*^ mice and in vivo selection of transgenic parasites with paromomycin. We engineered four transgenic *C. parvum* lines by replacing the parental *CpMetRS* locus with a transgene encoding wild-type *Cp*MetRS or *Cp*MetRSs with the D243E mutation, the T246I mutation, or both mutations in combination. The repair cassette was also designed to insert sequences for the enolase 3’ untranslated region (UTR) for transcriptional termination of *CpMetRS*, and a fused nanoluciferase (Nluc)-neomycin phosphotransferase (Neo) under control of a constitutively active enolase promoter located just after the *CpMetRS* stop codon (Fig. 2a). For each mutant parasite line, *CpMetRS* was recoded from base pairs 728 to 1686 to prevent homology directed repair without insertion of the selection marker. The recoded sequence also altered the gRNA recognition region to prevent repeated Cas9-mediated DNA cleavage (Fig. 2b). Each transgenic parasite line was generated without difficulty and each was passaged once in NOD SCID gamma mice under paromomycin selection to remove potential parental contaminants before subsequent experiments. PCR using flanking primers demonstrated insertion of the repair cassettes at the endogenous locus with amplification of the 1211 bp and 2945 bp bands predicted for the parental WT (Iowa) and transgenic parasite lines, respectively (Fig. 2c). Sanger sequencing confirmed the successful introduction of each desired mutation (Fig. 2d).

**Fig 2.**
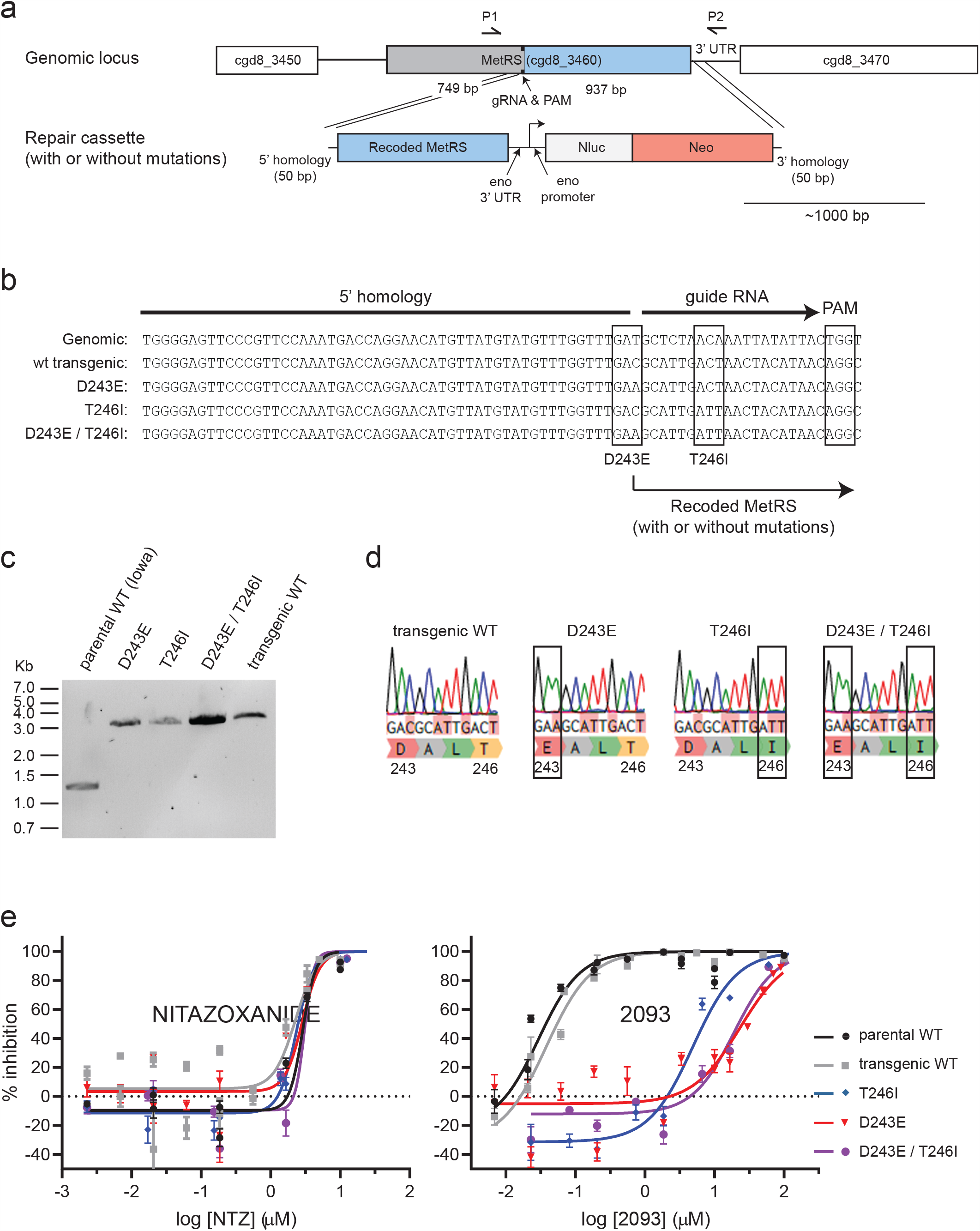
CRISPR/Cas9 engineered MetRS mutations confer *C. parvum* resistance to 2093. **(a)** Strategy used to modify the *C. parvum* MetRS genomic locus by CRISPR/Cas9 gene editing. Each DNA fragment is drawn to scale with an ∼1000 bp scale bar shown. **(b)** Aligned sequences showing the wild-type locus, wt transgenic locus, and intended mutations, along with the PAM CRISPR/Cas9 cut site and guide RNA. **(c)** PCR confirmation of the genomic insertion using primers P1 and P2 shown in (a). The anticipated PCR product sizes for the parental and transgenic strains were 1211 bp and 2945 bp, respectively. **(d)** Sanger sequencing chromatograms confirming introduction of each desired mutation. **(e)** Susceptibility of parental and transgenic *C. parvum* isolates to nitazoxanide (NTZ) (negative control) and **2093**. Data are combined from at least two biological replicate experiments per stain. Curves were calculated using all data points, but only every other dose is plotted in order to reduce clutter on the graphs.

We next purified oocysts for each transgenic *C. parvum* line from mouse feces, and the in vitro growth rates and drug susceptibility of each parasite line were assessed using a high content microscopy-based assay for parasite development (*39*). There were no significant differences in growth of the transgenic lines relative to Iowa strain *C. parvum*, as measured at 48 h post infection (Supplemental Fig. 3). The *Cp*MetRS mutations had no effect on *C. parvum* susceptibility to the unrelated growth inhibitor nitazoxanide (NTZ; negative control). On the other hand, the *MetRS* coding mutations reduced susceptibility to **2093** (Fig. 2e). The **2093** EC_50S_ for the D243E and T246I mutant MetRS *C. parvum* strains were 613- and 128-fold higher than those of the transgenic WT control, respectively (EC50_trangenic wt_ = 0.038 µM; EC50_D243E_ = 23.4 µM; EC50_T246I_ = 4.89 µM). **2093** resistance of the D243E / T246I double mutant was like that of the D243E single mutant. These data confirmed that the MetRS mutations selected in the calf model conferred *C. parvum* **2093** resistance, while not affecting susceptibility to nitazoxanide.

### Enzymatic properties of mutant *Cp*MetRS enzymes

The wild-type and two mutant *Cp*MetRS genes were cloned, overexpressed in *Escherichia coli*, and purified for enzymatic studies. Enzymatic activity was confirmed in the aminoacylation assay (*34*), which detects the esterification of radiolabeled methionine to the tRNA substrate. In order to measure the Michaelis-Menten constants for the enzymes, an ATP:PPi exchange assay was used, as previously explained (*40*). First, looking at wild-type enzyme, the *K*_*m*_ values for methionine and ATP were 31.3 and 1623 µM, respectively. These results are within a factor of ∼2 of previously published results (*34*). The *K*_*m*_ values for the two mutant enzymes were considerably lower than for the WT enzyme (Table 1), and the K_cat_/K_m_ values were higher, indicating greater catalytic efficiency. Next, the IC_50_ and *K*_*i*_ values of **2093** were determined for the three enzymes (Table 1). **2093** was extremely potent on the wild-type *Cp*MetRS with a *K*_*i*_ of 0.0017 nM. The IC_50S_ of **2093** against both mutant enzymes were greater than the top dilutions tested in two assays (>10 µM and >30 µM), indicating a >170-fold upward shift in the IC_50_ values.

**Table 1.** Kinetic parameters and sensitivity to **2093** inhibition of wild-type (WT) and mutant (T246I and D243E) *Cp*MetRS enzymes.

| Enzyme | Met titration |  |  | ATP titration |  |  | IC50 (nM)<br>for 2093 | Ki (nM) for<br>2093 |
| --- | --- | --- | --- | --- | --- | --- | --- | --- |
| | $K_m$<br>($\mu\text{M}$) | $k_{cat}$<br>($\text{S}^{-1}$) | $k_{cat}/K_m$($\mu\text{M}^{-1}\text{S}^{-1}$) | $K_m$<br>($\mu\text{M}$) | $k_{cat}$<br>($\text{S}^{-1}$) | $k_{cat}/K_m$($\mu\text{M}^{-1}\text{S}^{-1}$) | | |
| WT | 31.3 | 30.6 | 0.98 | 1623 | 99.0 | 0.064 | $174.1 \pm 23.3$<br>(n=2) | $0.0017 \pm 0.0003$ (n=2) |
| T246I | 2.93 | 116.8 | 39.8 | 410.3 | 111.2 | 0.271 | >10,000,<br>>30,000 | >31.8, >99.9 |
| D243E | 6.94 | 158.6 | 22.8 | 890.6 | 138.9 | 0.156 | >10,000,<br>>30,000 | >33.3, >99.9 |

### Mechanism of compound 2093 resistance

We previously solved multiple crystal structures of the *Trypanosoma brucei (Tb)* MetRS with inhibitors bound, including with **2093** (PDB: 6CML), which demonstrated that they inhibit enzymatic activity by competing with substrate binding (*41-44*). The *Cp*MetRS and *Tb*MetRS are 76% identical (19 of 25 amino acids) within the inhibitor binding pocket (*34*), which presented the opportunity to further examine the mechanism of *Cp*MetRS resistance to **2093**. We generated a *Cp*MetRS model structure based on the *T. brucei* structures using I-TASSER (*45*), and then docked compound **2093** into the model (Fig. 3). The predicted binding mode of **2093** to *C. parvum* MetRS shows the 2-chloro,4-methoxy-benzyl positioned in the enlarged methionine substrate pocket and the 5-Cl-imidazo[4,5b]pyridine in the “auxiliary pocket” formed upon inhibitor binding *(41, 42)*(also explained in Materials and Methods). D243 forms a salt bridge with H63. This salt bridge is at 3.8 Å from both the C6 and C7 of the imidazo[4,5-b]pyridine bicyclic system. We hypothesize that the D243E mutation disrupts the side wall of the auxiliary pocket by virtue of its larger size, which would lead to less favorable binding of the inhibitor. T246 is within van der Waals distance of the F242 phenyl ring. The resistance mutation T246I probably leads to a clash with the phenyl ring and thereby causes local unfolding of the ***α***-helix it belongs to, thereby disrupting the back wall of the auxiliary pocket.

**Fig 3.**
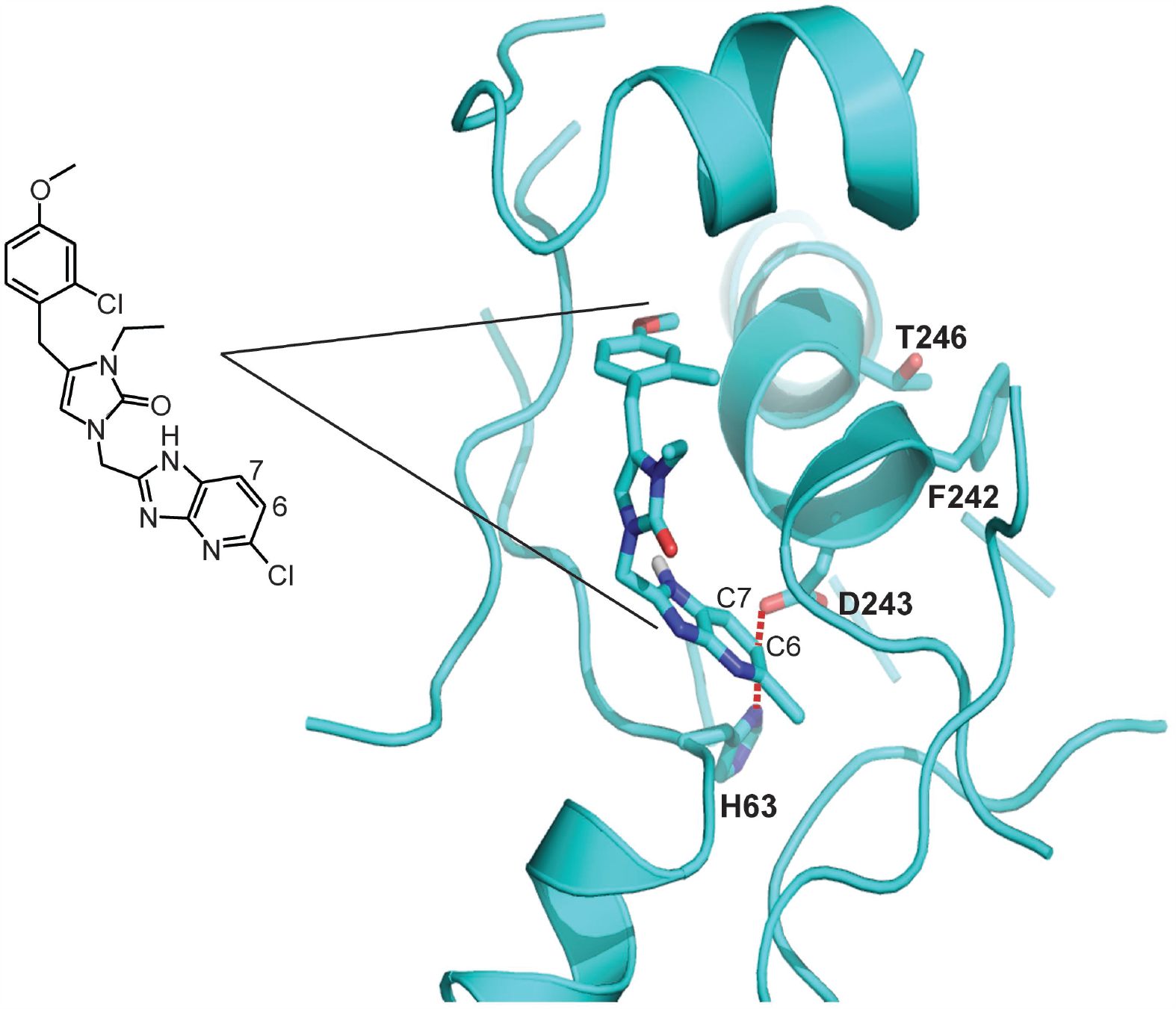
2093 docked into a comparative model of *Cp*MetRS. **2093** is coded with C: cyan, N: blue, O: red. The resistance mutations D243 and T246 are labeled. D243 forms a salt bridge with H63. This salt bridge is at 3.8 Å from both the C6 and C7 of the imidazo[4,5-b]pyridine bicyclic system. T246 is within van der Waals distance of the F242 phenyl ring. **2093** is also shown as a stick diagram for reference.

## Discussion

This study is important because it highlights the likelihood of *Cryptosporidium* drug resistance, which has thus far been neglected during anticryptosporidial drug development. Resistance to the *Cp*MetRS inhibitor **2093** arose after four days of drug exposure in the neonatal dairy calf model despite plasma and fecal concentrations well above the in vitro EC_90_ for **2093** (Supplemental Fig. 1b). Two amino acid altering SNVs in *CpMetRS* emerged independently, a D243E mutation and a T246I mutation. CRISPR/Cas9 genome editing showed that these mutations confer high-level *C. parvum* resistance to **2093**, while not significantly affecting parasite growth or susceptibility to nitazoxanide. In studies using recombinant enzymes, the D243E and T246I mutations shift the **2093** IC_50_ by >170-fold. Modeling based on solved *Tb*MetRS crystal structures suggests that the resistance mutations reposition nearby hydrophobic residues, interfering with compound binding while minimally impacting substrate binding. Our data confirm that the anticryptosporidial activity of **2093** and its analogs occurs via competitive inhibition of *Cp*MetRS, and, more importantly, demonstrate rapid emergence of drug resistant *C. parvum* mutants.

Drug resistance in our study resulted from two coding single nucleotide alterations in the *CpMetRS*. It is uncertain if parasites harboring these mutations were pre-existing in the large population of parasites used to initiate infection or arose by spontaneous mutation in the presence of selective drug pressure. Suggesting a mechanism of spontaneous mutation, the *MetRS* SNVs identified here are not reported in the 53 *Cryptosporidium* genomes currently available in CryptoDB (https://cryptodb.org/cryptodb/). There are also no MetRS inhibitors in agricultural use to which the *C. parvum* isolate used in our calf study may have been previously exposed.

Evolution of antimicrobial resistance through spontaneous mutation is favored by the presence of large numbers of organisms, which could provide an explanation for outgrowth of resistant strains in the calf model despite its previously documented efficacy in murine models. For example, Gram-negative bacterial resistance with specific target mutations developed in three of fourteen people after only one day of treatment in a phase II human trial of the LeuRS inhibitor GSK2251052 (AN3365) for complicated urinary tract infections (*27*). This occurred despite an inability to select for GSK2251052 resistance in mice, presumably due to lower numbers of organisms in the mouse model. In our calf study, peak oocyst shedding in each control animal ranged from 10^7^ to 10^8^ oocysts per gram of feces. This number is consistent with previous studies of *C. parvum* infected calves (*19, 37*) and, based on the approximate volume of feces produced by a calf with cryptosporidiosis, on the order of 10^10^ to 10^11^ oocysts are shed during a typical ten day experiment using this model (*19, 37, 46*). If we look to another apicomplexan parasite, *Plasmodium falciparum*, the estimated number of *C. parvum* organisms in the calf model is in-line with the burden of organisms during acute falciparum malaria (estimated at 10^8^ to 10^12^ (*47*)), a scenario in which treatment failure due to newly derived resistance is well-documented during treatment with a single drug, e.g. with atovaquone (*48*). Up to 1 in 10^5^ malaria parasites develop resistance to atovaquone (*49*), virtually guaranteeing the development of resistance on monotherapy. We also note that the simplistic view of microbial evolution described above ignores the potential for sexual recombination within the human host, a life cycle feature of *Cryptosporidium* species that distinguishes it from *Plasmodium* and provides an additional mechanism to generate genetic diversity in the face of selective pressure. A key unknown for cryptosporidiosis is the number of organisms during a human infection, which may of course differ for different host groups (e.g. parasite numbers might be greater in AIDS patients than in immunocompetent hosts).

The types and number of mutations required to confer antimicrobial resistance is a specific characteristic of different drug classes and targets, and together with fitness costs resulting from resistance mutations determines the propensity for emergence of resistance to a given drug (*22*). For example, *S. aureus* resistance to rifampin typically results from one of several single amino acid mutations in the RNA polymerase β subunit (rpoB) that emerge readily (*50, 51*); as a result, rifampin is not useful as monotherapy for *S. aureus* infections. Studies with a first-generation MetRS inhibitor, REP8839, showed resistance frequency rates for *S. aureus* in the same range as rifampin (10^−7^ to 10^−8^ over a 48 hour exposure) (*52*), suggesting a low barrier to resistance. The first-step REP8839 *Staphylococcus* resistance SNVs occurred in close proximity to the corresponding *CpMetRS* SNVs identified in this study *(52)*, and the effects of these SNVs on enzyme kinetics with the natural substrates (methionine and ATP) were small for *Staphylococcus* as well as for *Cryptosporidium*. These findings raise concerns about developing MetRS inhibitors or other enzyme inhibitors that are vulnerable to resistance by subtle mutations in *Cryptosporidium*’s genome. The use of MetRS inhibitors as part of combination therapy may still be a consideration given their in vivo activity. Furthermore, later-generation MetRS inhibitors have been developed with much lower resistance frequency rates to *S. aureus* (on the order of 10^−10^) (unpublished, F. Buckner and E. Fan), suggesting the potential for discovering MetRS inhibitors with higher barriers to resistance in *Cryptosporidium*.

There are several limitations to our study, which was initially designed as a drug efficacy trial and not intended to be a comprehensive study of *Cryptosporidium* drug resistance. Key factors likely to affect emergence of MetRS inhibitor resistance and the general likelihood of *Cryptosporidium* resistance developing via target mutations remain unknown, including knowledge of *Cryptosporidium’s* spontaneous mutation rate, the impact of sexual recombination in the presence of drug pressure, and the effect of target mutations on parasite fitness. Furthermore, our data only address the *Cp*MetRS target mutations that spontaneously emerged in the calf model. Alternative mechanisms of drug resistance such as altered expression of MDR efflux pumps or other compensatory alterations in gene expression remain completely unexplored. Such resistance mechanisms may operate independently during evolution of drug resistance or may mediate drug tolerance that enables parasite persistence while more resistant mutants evolve. We are currently focused on developing practical methods (i.e. not requiring dairy calves) to more completely characterize the evolution of *Cryptosporidium* MetRS inhibitor resistance and resistance to other classes of anticryptosporidials in the developmental pipeline with the goal of identifying suitable drugs or drug combinations. Methods to determine the relative propensities of different anticryptosporidials to induce resistance might guide preferential investment in compounds with high barriers to resistance. Furthermore, anticryptosporidial therapies based on combinations of drugs or drug-potentiator combinations may be identified to mitigate emergence of resistance, as is now standard for tuberculosis (*53*), HIV (*54*), and anti-cancer therapies (*22, 55*).

In summary, these data demonstrate spontaneous emergence of drug resistant *C. parvum* during a dairy calf clinical efficacy trial of the *Cp*MetRS inhibitor **2093**. Resistance was mediated by *CpMetRS* mutations coding altered amino acids that appear to reduce compound binding with minimal effect on substrate binding. Rapid evolution of resistance was likely favored by the large burden of *C. parvum* in the dairy calf model; it remains unknown if other mechanisms mediating tolerance or resistance occurred that might have enabled parasite persistence and thereby facilitated emergence of highly resistant *CpMetRS* mutants. Our experience heightens concern about the inevitability of *Cryptosporidium* drug resistance. The anticryptosporidial field is poised for introduction of improved treatments in the next several years. With the goal of understanding how to best deploy these new treatments, we are now developing methods to systematically study anticryptosporidial drug resistance and identify approaches to impede its occurrence.

## Materials and Methods

### Neonatal calf model of cryptosporidiosis

Animal studies were approved by the University of Vermont (UVM) Institutional Animal Care and Use Committee (IACUC). The University of Vermont has an Animal Welfare Assurance with the Office of Laboratory Animal Welfare (OLAW) of the National Institutes of Health and is registered as a Research Institution by the United States Department of Agriculture. The work was performed in compliance with the Guide for the Care and Use of Laboratory Animals and with the Animal Welfare Act and its associated regulations (USDA-APHIS “Blue Book” (www.aphis.usda.gov/animal-welfare)).

Holstein bull calves were acquired at birth from Green Mountain Dairy (Sheldon, VT), given synthetic colostrum with 200 g of IgG (Land O’Lakes, Ardent Hills, MO) and bovine coronavirus and *Escherichia coli* antibodies (First Defense Bolus, Immuncell Corporation, Portland, ME) within two hours of birth, and transported to UVM. Uninfected animals were group-housed and infected at 24-48 hours of age during an interruption in bottle feeding by oral administration of ∼5×10^7^ viable *C. parvum* Iowa isolate oocysts (Bunch Grass Farms, Deary, ID) suspended in 10 mL of deionized water. Animals were moved to individual raised pens immediately after infection. Animals with severe diarrhea and other symptoms were supported with oral electrolytes or intravenous fluids, and flunixin meglumine (Banamine, Merck) as needed. **2093** was suspended for dosing in 10 mL of 60% Phosal 53 MCT Lipoid, 30% PEG400, and 10% ethanol by volume. Doses were prepared fresh each day from solid **2093** and squirted into the calves’ mouths during interruptions in bottle feeding. For PK studies, fecal samples were collected at the indicated times by manual anal stimulation. For treatment efficacy studies, daily fecal samples were obtained from collection bins located under each pen. Fecal samples used for parasite quantification were dried at 90 °C until a stable weight was reached, and *C. parvum* abundance per gram of fecal dry matter was measured using a previously validated qPCR assay (primers listed in Supplemental Table 1) (*56*). The lower limit of detection for this qPCR assay is ∼100 oocysts/g of dried feces.

### Pharmacologic methods

Serum samples were analyzed by liquid chromatography-tandem mass spectrometry (LC/MS/MS), using compound spiked into control serum as a standard. Fecal compound **2093** was measured by homogenizing feces in PBS (0.1 g/mL) in a polypropylene tube and then further dilution prior to addition of an internal standard (enalapril) and acetonitrile protein precipitation. The supernatant was transferred to a fresh tube and dried using a speed vac. Samples were then resuspended and analyzed by LC/MS/MS.

### PCR analysis of *CpMetRS* in shed oocysts

Oocyst DNA was prepared from fecal samples (as described above). The *CpMetRS* gene was Sanger sequenced on day 3 (3 vehicle calves, 3 treatment calves) and on day 9 (single surviving vehicle calf and 2 treatment calves which showed a relapse). The Day 9 samples from the treatment group which showed point mutations were then amplified from genomic DNA, gel purified using the QiaQuick Gel Extraction Kit, (Qiagen, NL), cloned into the AVA0421 plasmid (*57*), and sequence verified.

### Cell-based *C. parvum* growth inhibition assay

A previously established high-content microscopy assay was used to measure activity of **2093** against the parental and transgenic *C. parvum* strains when grown in the human colon cancer cell line HCT-8 (ATCC CCL-244) (*39*). HCT-8 cells were grown in RPMI 1640 medium (Invitrogen) supplemented with 10% heat-inactivated fetal bovine serum (Sigma-Aldrich), 120 U/mL penicillin, and 120 µg/mL streptomycin (ATCC) at 37°C to near confluence in clear-bottomed 384-well plates in 50 µL per well. *C. parvum* oocysts were induced to excyst by treatment with 10 mM hydrochloric acid (10 min at 37°C) and then 2 mM sodium taurocholate in PBS (10 min at 16°C), washed and resuspended in the above medium, and then added to cell monolayers at a concentration of 5.5×10^3^ per well. Compounds were added 3 h after infection, and assay plates were incubated for 48 h post-infection at 37°C under 5% CO_2_. Wells were then washed three times with PBS containing 111 mM _D_-galactose, fixed with 4% formaldehyde in PBS for 15 min at room temperature, permeabilized with 0.25% Triton X-100 (10 min at 37°C), washed three times with PBS with 0.1% Tween 20, and blocked with 4% bovine serum albumin (BSA) in PBS for 2 h at 37°C. Parasitophorous vacuoles were stained with 1.33 µg/mL fluorescein-labeled *Vicia villosa* lectin (Vector Laboratories) diluted in 1% BSA in PBS with 0.1% Tween 20 (1 h at 37°C), followed by the addition of Hoechst 33258 (AnaSpec) at a final concentration of 0.09 mM diluted in water (15 min at 37°C). Wells were then washed five times with PBS containing 0.1% Tween 20. A Nikon Eclipse TE2000 epifluorescence microscope with an EXi Blue fluorescence microscopy camera (Qimaging, Canada) with a 20× objective (0.45 numerical aperture). Nucleus and parasite images were exported separately as .tif files and were analyzed using macros developed on the ImageJ platform (National Institutes of Health) to determine the numbers of parasites and host cells (*39*).

### Generation of transgenic *C. parvum* by CRISPR/Cas9 genome editing

Transgenic *C. parvum* were generated using CRISPR/Cas9 genome editing and a previously reported method employing in vitro electroporation of excysted sporozoites followed by infection of *ifn-γ* ^*-/-*^ mice and paromomycin selection (*38*). Mouse experiments for generation of transgenic *C. parvum* were approved by the University of Vermont Institutional Animal Care and Use Committee (IACUC).

To generate the Cas9/guide RNA plasmid, the *CpMetRS* guide sequence was cloned into the *Bbs*I restriction site of the plasmid Aldo-Cas9-ribo. For the repair construct, a synthetic 999 bp fragment of the recodonized version of the *CpMetRS* (starting from amino acid number 247) along with the 3’UTR sequence of the enolase gene (cgd5_1960) was purchased (GenScript, NJ, USA). The synthetic construct was PCR amplified to introduce desired mutations and the 5’ homology region was introduced as an overhang in the forward primer. The enolase promoter-nluc-neomycin cassette was amplified from the vector Cplic3HAENNE provided by the Striepen Lab (Univ. of Pennsylvania) and the 3’ homology region was added at the end of this construct as a primer overhang. A 40 bp sequence homology was introduced in the two above mentioned fragments during the PCR and they were ligated with the NEBuilder HiFi DNA Assembly Master Mix (New England Biolabs; catalog number E5520S). The whole cassette was then amplified with the end primer set. Each re-codonized construct was cloned into pCR™4Blunt-TOPO vector (Invitrogen) and sequence verified. To transfect *C. parvum*, the repair cassette for each mutant was PCR amplified from this vector using the end primer set. All PCR primers are included in Supplemental Table 1.

*Cryptosporidium parvum* Iowa strain oocysts were purchased from Bunch Grass Farm (Deary, ID), excysted by treatment with 10 mM hydrochloric acid (10 min at 37°C) and then 2 mM sodium taurocholate in PBS (10 min at 16°C), and electroporated with Cas9/guide plasmid and repair DNA template using a Lonza Nucleofector 4D electroporator and the previously reported protocol (*38*). C57BL/6 *ifn-γ* ^*-/-*^ mice (Jackson Laboratory) aged 4-6 weeks (n=2) were infected by injection of transfected sporozoites into an externalized loop of small intestine, and transgenic parasites were selected for by inclusion of paromomycin (Gemini #400-155P, 20 g/liter) in the mouse drinking water. Fecal samples were collected from cages and luminescence measurements were performed as described previously (*38*). Mouse feces containing transgenic parasites were then passaged by infection of NOD SCID gamma mice (Jackson Laboratory), which become chronically infected with *C. parvum (15)*. Oocysts were purified from these mice for subsequent experiments.

Primers complementary to the 5’ homologous *CpMetRS* region and the *CpMetRS* 3’UTR (primers P1 and P2 in Supplemental Table 1) were used for PCR on fecal DNA from the infected NOD-SCID gamma mice to confirm the correct 5’ and 3’ integration events following homologous recombination. For this, total fecal DNA was purified using an E.Z.N.A. Stool DNA Kit (Omega Bio-Tek), according to the manufacturer’s pathogen detection protocol except for inclusion of six freeze-thaw cycles in kit lysis buffer as a first step. Sanger sequencing was used to confirm incorporation of the desired mutations, and parasites were purified for use in dose-response assays by sucrose flotation followed by cesium chloride purification using a previously described protocol (*58*). Mouse fecal pellets were collected every two hours for oocyst purification, which enhanced parasite viability by preventing desiccation.

### Recombinant *Cp*MetRS expression and in vitro enzymatic studies

The expression and purification of the *Cp*MetRS proteins was performed as previously described (*57*) with a few changes as follows. PA0.5G non-induction media was prepared with 10× metals mix and 100 µg/ml each of L-methionine and 17 amino acids. Cultures were then incubated overnight at 37°C. ZYP-5052 media was prepared with ampicillin 100 µg/ml, carbenicillin 100 µg/ml and chloramphenicol 34 µg/ml and incubated for 24-48 hours at room temperature. The protein/Ni^2+^ mixture was incubated overnight at 4°C. Wash buffer contained 50mM imidazole and 8 washes were performed, while elution buffer had 250 mM imidazole and eluted proteins were concentrated down to ∼100 µl using Amicon Ultra 0.5 mL Centrifugal Filters (Amicon, Darmstadt, Germany).

Enzyme kinetics and **2093** *K*_*i*_ values for *Cp*MetRS WT, *Cp*MetRS 74-9 (T246I) and *Cp*MetRS 76-9 (D243E) were measured using an ATP:PPi exchange assay performed at room temperature (*40*). For *Ki* measurements, **2093** was preincubated for 5 min in a 96 well-plate with 30 nM *Cp*MetRS, 50 mM L-methionine, 2.5mM NaPPi, 3 µCi of [32P]tetrasodium pyrophosphate (NEX019005MC; Perkin-Elmer), 2.5 mM DTT, 0.1 mg/ml BSA, 0.2mM spermine, 25 mM HEPES-KOH (pH 7.9), 10 mM MgCl_2_, 50 mM KCl, and 2% DMSO.

Reactions were started with addition of 2.5mM ATP; after 20 min, 5 µl of the reaction mixture (in duplicate) was quenched into a MultiScreenHTS Durapore 96-well filter plate (MSHVN4B50; Millipore Sigma) containing 200 µl of 10% charcoal in 0.5% HCl and 50 µl of wash buffer (1 M HCl with 200 mM NaPPi). The filter plates were washed three times with 200 µl of wash buffer on a vacuum manifold. The plates were dried for 1 hour at RT, and then 25 µl of scintillation cocktail was added. Counts per minute (CPM) were quantified on a MicroBeta2 scintillation counter (Perkin-Elmer). Percent inhibition was calculated by subtracting the background wells (containing all assay reagents except ATP, L-Methionine and compound) and comparing this value to the high control wells (containing all assay reagents without compound). The IC_50_ values were calculated by nonlinear regression methods using the Collaborative Drug Database (Burlingame, CA; www.collaborativedrug.com). K_i_ values were calculated from the IC_50S_ that were shifted above the enzyme concentration (30 nM) using the Cheng-Prusoff equation as follows: IC_50_ = (1 + [Met]/K_m_ ^Met^)(1 + K_m_ ^ATP^/[ATP])Ki (*40*). The K_m_ values for L-methionine was determined using the assay conditions described above for each *Cp*MetRS enzyme (30 nM) without compounds and with 5 µCi of [^32^P]tetrasodium pyrophosphate, 2.5 mM ATP, and 2.5 mM NaPPi, while titrating L-methionine. The reaction was quenched as described above at time intervals of 0, 4, 8, 12, 16, and 20 min. Similarly, the K_m_ values for ATP were measured by using 1 mM L-methionine and 2.5 mM NaPPi while titrating ATP. The K_m_ and V_max_ values were calculated using GraphPad Prism (v6.0). The k_cat_ is equal to the V_max_ divided by the enzyme concentration (30 nM). All IC_50_ and K_m_ determination assays were performed twice to ensure reproducibility.

### Comparative modeling of *Cp*MetRS and docking of compound 2093

A 3D model for UNPROT amino acid sequence Q5CVN0 was constructed with I-TASSER (*59*). A user specified template was provided using the crystal structure of *Trypanosoma brucei* MetRS in complex with inhibitor **1312** (2-((3-((3,5-dichlorobenzyl)amino)propyl)amino)quinolin-4(1H)-one, PDB: 4eg5, chain B) to ensure a conformation of the protein that possesses the “auxiliary pocket” seen with dozens of inhibitors of that type, not only with *T*.*brucei* MetRS (*41, 42*) but also the congeners of *Brucella melitensis (60) and Staphylococcus aureus* (unpublished, PDB: 4qrd, 4qre). The overall sequence identity between template and target was 37%, the similarity 53%; the sequence identity in the expected binding site is 76%). Because I-TASSER does not take into account the presence of the ligand manual rotamer adjustment for five amino acid residues in the ligand binding site was necessary (residues Y24, H63, T185, T242, R221) so that they were the same as in the *T*.*brucei* MetRS complex.

Subsequently **2093** was docked in the 3D model of *C. parvu*m MetRS with the *Monte Carlo* algorithm implemented in QXP+ (version 2016) (*61*), allowing for full flexibility of the ligand and the amino acid residues in direct contact.

### Data analysis, statistical methods, and figure preparation

Data were analyzed using GraphPad Prism version 7.01, and graphs were then exported as .eps files. The area under the curve (AUC) for oocyst shedding in calf feces was calculated for each animal using a plot of Log_10_ transformed fecal oocyst shedding per gram of fecal dry matter vs. time from days two to six with a baseline of 2 (Log_10_=100). The P value for oocyst shedding was determined using an unpaired 2-tailed t test. Oocyst growth index data were assessed for normality using the D’Agostino & Pearson normality test. Because the measured growth indices for the D243E/T246I double mutant cell line failed the normality test, the mean growth indices of each cell line were compared to that of Iowa strain parasites using the non-parametric Kruskal-Wallis and Dunn’s multiple comparisons tests. Dose response curves and half-maximal effective concentrations (EC_50S_) were determined by nonlinear regression using the equation for log (inhibitor) vs. response curve with variable slope with the top response constrained to 100%. Figures were prepared using Adobe Illustrator CS5.

## Acknowledgements

We are grateful to Colin McMartin for providing us with the QXP+ software.

## Author contributions

MMH: Conceptualization, methodology, validation, formal analysis, visualization, writing (original draft, review, and editing); ES: Conceptualization, methodology, formal analysis; RKMC: Conceptualization, writing (review and editing), funding acquisition; JRG: Methodology, validation, formal analysis, writing (original draft, review, and editing), visualization; ELdH: Conceptualization, writing (review and editing), funding acquisition; PM: Methodology, validation, writing (review and editing); AM: Methodology, validation, formal analysis, writing (original draft); RMR: Methodology, validation, formal analysis; JET: Conceptualization, methodology, writing (review and editing); CLMJV: Methodology, writing (original draft, review and editing), visualization; AS: Conceptualization, methodology, writing (review and editing); ZZ: Methodology; DMO: Methodology; formal analysis; DWG: Methodology; formal analysis; EF: Writing (review and editing), supervision, funding acquisition; FSB: Conceptualization, methodology, writing (original draft, review and editing), supervision; funding acquisition; CDH: Conceptualization, methodology, formal analysis, visualization, writing (original draft, review and editing), supervision, funding acquisition.

## Data and materials availability

All data reported in this study are presented in the paper and/or supplementary materials. The plasmids and genetically modified *C. parvum* strains are available from CDH under a material transfer agreement (MTA) with the University of Vermont Larner College of Medicine. The plasmids for expression of recombinant *Cp*MetRS proteins are available from FSB under an MTA with the University of Washington.

## Supplementary Materials

**Supplemental Fig. 1. Serum and fecal 2093 pharmacokinetics in uninfected and *C. parvum* infected bull calves.** Serum and fecal **2093** concentrations were measured using LC-MS/MS. A horizontal line is drawn for reference on each graph at three times the EC_90_ of **2093** measured in HCT-8 cells, which is the concentration previously shown to be needed to maximize the rate of parasite elimination (*34*). **(a)** Single-oral dose PK study. Blood and fecal samples were collected at 0, 0.5, 1, 2, 4, 6, 8, 12, and 24 h after administering a single oral 10 mg per kg dose of **2093**. Each color-coded line indicates an individual animal (n=2). **(b)** Serum and fecal **2093** concentrations in *C. parvum* infected calves. Blood and fecal samples were collected at 5, 10, 24, 34, 72, and 168 h after beginning **2093** treatment at a dose of 15 mg per kg twice daily. Each color-coded line indicates an individual animal (n=3 except for missed samples post 72 h).

**Supplemental Fig. 2. DNA sequencing results for *CpMetRS* shed during 2093 treatment of *C. parvum* infected dairy calves. (a)** Predicted *Cp*MetRS amino acid sequences obtained prior to treatment and on experiment day 9. The calf number and day of experiment is indicated. The sequence from calf 74 day 1 (prior to treatment) is identical to the *Cp*MetRS in the cryptoDB genome database. The box highlights the predicted amino acid mutations in the *Cp*MetRS of parasites shed by calves 74 and 76 on day 9. **(b)** Sanger sequencing chromatograms and DNA sequences for oocysts shed by calves 74 and 76 on day 9.

**Supplemental Fig. 3. In vitro growth of transgenic *C. parvum* lines relative to Iowa strain *C. parvum*.** HCT-8 cell monolayers were infected with each parasite line for 48 h, followed by enumeration of parasites and host cell nuclei by high content microscopy (*39*). Data are shown as a growth index, calculated as the percent of host cells infected normalized to the mean percent of host cells infected by Iowa strain parasites on the same assay plate (n=8 culture wells per *C. parvum* line; no significant differences by Kruskal-Wallis and Dunn’s multiple comparisons tests).

**Supplemental Table 1. PCR primers**.

