## Supplemental Figure 1 for "Spontaneous selection of *Cryptosporidium* drug resistance in a calf model of infection"

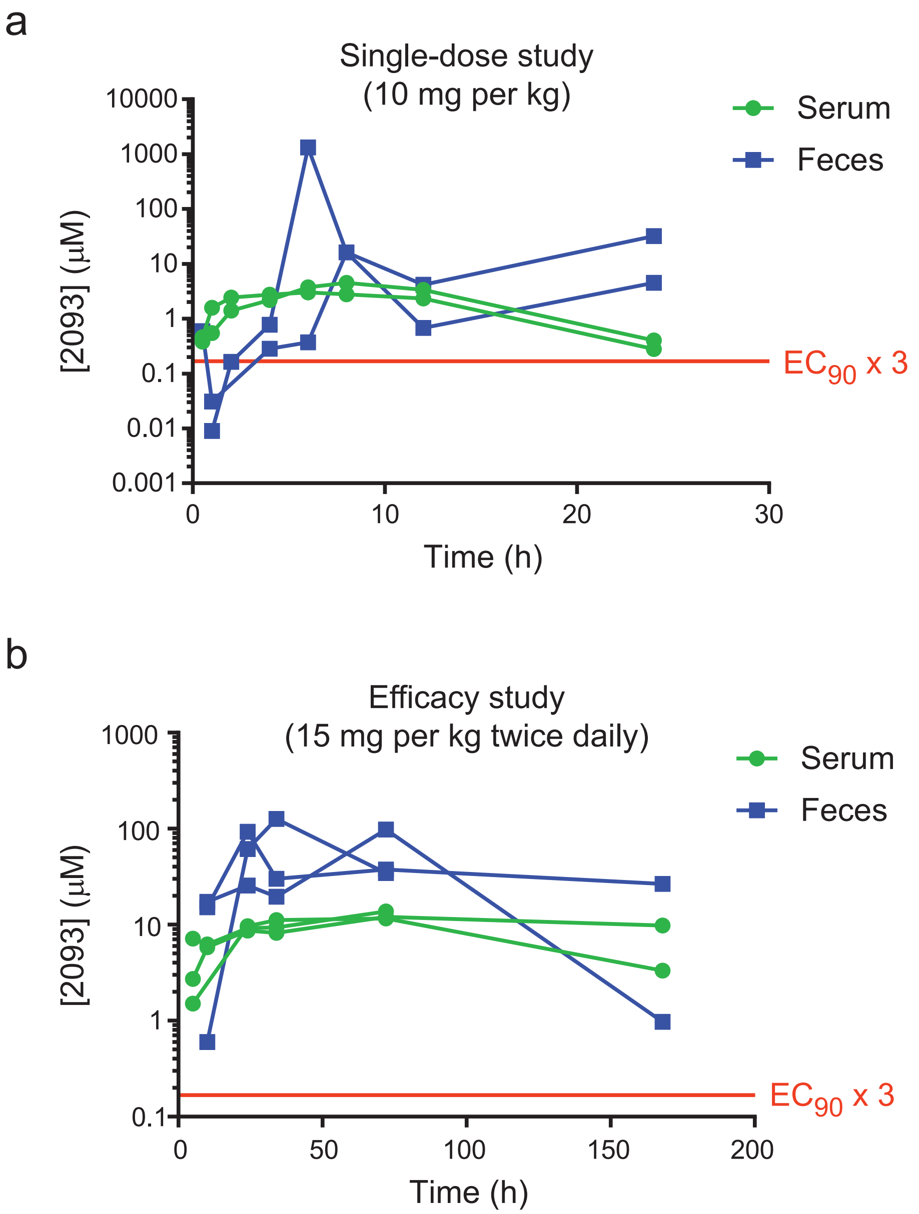


**Supplemental Figure 1: Serum and Fecal compound 2093 pharmacokinetics in uninfected and *C. parvum* infected bull calves.** Serum and fecal compound 2093 concentrations were measured using LC-MS/MS. A horizontal line is drawn for reference on each graph at three times the $EC_{90}$ of compound 2093 measured in HCT-8 cells, which is the concentration previously shown to be needed to maximize the rate of parasite elimination *(34)*. **(a)** Single-oral dose PK study. Blood and fecal samples were collected at 0, 0.5, 1, 2, 4, 6, 8, 12, and 24 h after administering a single oral 10 mg per kg dose of compound 2093. Each color-coded line indicates an individual animal (n=2). **(b)** Serum and fecal compound 2093 concentrations in *C. parvum* infected calves. Blood and fecal samples were collected at 5, 10, 24, 34, 72, and 168 h after beginning compound 2093 treatment at a dose of 15 mg per kg twice daily. Each color-coded line indicates an individual animal (n=3 except for missing samples post 72 h).
