## Supplemental Figure 2 for "Spontaneous selection of *Cryptosporidium* drug resistance in a calf model of infection"

**a**

```
Calf74_day1      MTKINTTYRNSSGKYYYFTTAINYTNGTPHMGHAYEIVTSDILVRIARLFGFNTRFQTGT  60
Calf74_day9      MTKINTTYRNSSGKYYYFTTAINYTNGTPHMGHAYEIVTSDILVRIARLFGFNTRFQTGT  60
Calf76_day9      MTKINTTYRNSSGKYYYFTTAINYTNGTPHMGHAYEIVTSDILVRIARLFGFNTRFQTGT  60
                 ************************************************************

Calf74_day1      DEHGQKIAATAEKKGISPKELCDENVKQFKNMNEKLQISTDRFIRTTDADHYESCKELWR  120
Calf74_day9      DEHGQKIAATAEKKGISPKELCDENVKQFKNMNEKLQISTDRFIRTTDADHYESCKELWR  120
Calf76_day9      DEHGQKIAATAEKKGISPKELCDENVKQFKNMNEKLQISTDRFIRTTDADHYESCKELWR  120
                 ************************************************************

Calf74_day1      RCEAKGDIYLGKYNGWYNVREESFMTELEAKMLNYTDPISGLPLTKMEEPSYFFRITNYL  180
Calf74_day9      RCEAKGDIYLGKYNGWYNVREESFMTELEAKMLNYTDPISGLPLTKMEEPSYFFRITNYL  180
Calf76_day9      RCEAKGDIYLGKYNGWYNVREESFMTELEAKMLNYTDPISGLPLTKMEEPSYFFRITNYL  180
                 ************************************************************

Calf74_day1      ESIKNHIIDNPEFIQPESSRESILARLEALGSDSEDLSISRASFSWGVPVPNDQEHVMYV  240
Calf74_day9      ESIKNHIIDNPEFIQPESSRESILARLEALGSDSEDLSISRASFSWGVPVPNDQEHVMYV  240
Calf76_day9      ESIKNHIIDNPEFIQPESSRESILARLEALGSDSEDLSISRASFSWGVPVPNDQEHVMYV  240
                 ************************************************************

Calf74_day1      WFDALTNYITGLKWADFDNDLEENSLFNKYWENTVHIVGKDITWFHSVIWPAMLLSVGLS  300
Calf74_day9      WFDALINYITGLKWADFDNDLEENSLFNKYWENTVHIVGKDITWFHSVIWPAMLLSVGLS  300
Calf76_day9      WFEALTNYITGLKWADFDNDLEENSLFNKYWENTVHIVGKDITWFHSVIWPAMLLSVGLS  300
                 **:** ******************************************************

Calf74_day1      LPKTIFAHGFVTAPDGKKMSKSLGNVVDPFEQIDKIGSDPFRYYLAREGRFGNDIKYQPS  360
Calf74_day9      LPKTIFAHGFVTAPDGKKMSKSLGNVVDPFEQIDKIGSDPFRYYLAREGRFGNDIKYQPS  360
Calf76_day9      LPKTIFAHGFVTAPDGKKMSKSLGNVVDPFEQIDKIGSDPFRYYLAREGRFGNDIKYQPS  360
                 ************************************************************

Calf74_day1      SVIDYNNGELADTYGNLISRITNLTHKFCQGVAPSLSQSLLLDKPVDIDTVFEEYYLAWN  420
Calf74_day9      SVIDYNNGELADTYGNLISRITNLTHKFCQGVAPSLSQSLLLDKPVDIDTVFEEYYLAWN  420
Calf76_day9      SVIDYNNGELADTYGNLISRITNLTHKFCQGVAPSLSQSLLLDKPVDIDTVFEEYYLAWN  420
                 ************************************************************

Calf74_day1      CFRIDECIQIAMDLSRNINKFLTDHAPWNKDNGKSDEERIEIIRIVLEGSYYISILLSPF  480
Calf74_day9      CFRIDECIQIAMDLSRNINKFLTDHAPWNKDNGKSDEERIEIIRIVLEGSYYISILLSPF  480
Calf76_day9      CFRIDECIQIAMDLSRNINKFLTDHAPWNKDNGKSDEERIEIIRIVLEGSYYISILLSPF  480
                 ************************************************************

Calf74_day1      ITLACKEVFNRLGTPEKIIPEVDINYNNIKPGTPILIGEPLFKRIVNKEEKTFEELKAEK  540
Calf74_day9      ITLACKEVFNRLGTPEKIIPEVDINYNNIKPGTPILIGEPLFKRIVNKEEKTFEELKAEK  540
Calf76_day9      ITLACKEVFNRLGTPEKIIPEVDINYNNIKPGTPILIGEPLFKRIVNKEEKTFEELKAEK  540
                 ************************************************************

Calf74_day1      REREKQRRSGNKKLSETKDKGK 562
Calf74_day9      REREKQRRSGNKKLSETKDKGK 562
Calf76_day9      REREKQRRSGNKKLSETKDKGK 562
                 **********************
```

**b**

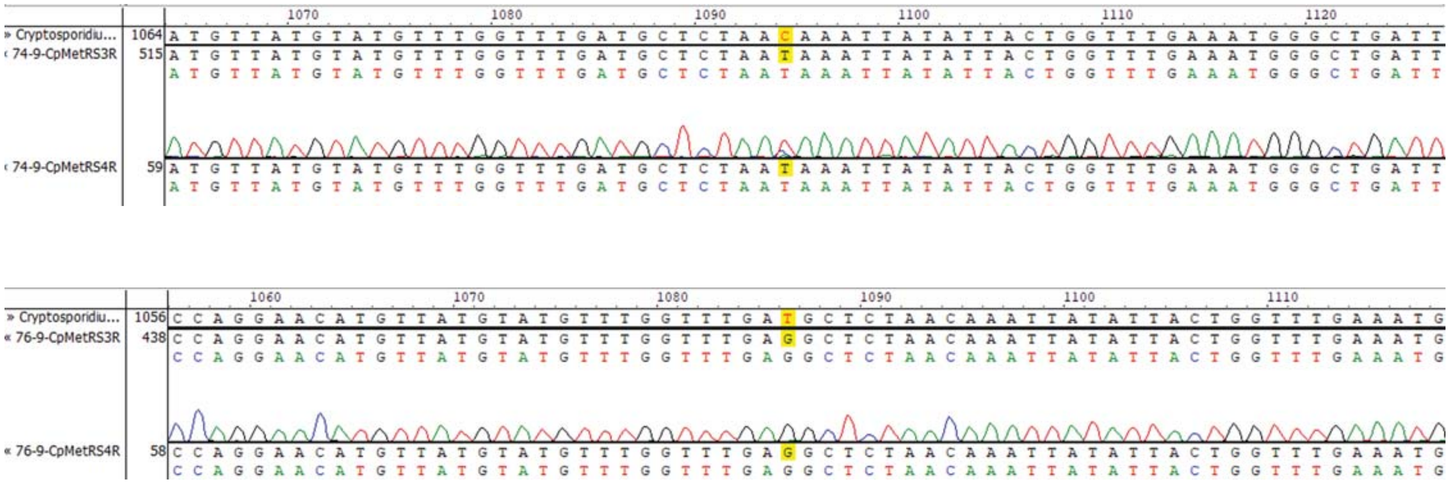


**Supplemental Fig. 2. DNA sequencing results for *CpMetRS* shed during 2093 treatment of *C. parvum* infected dairy calves. (a)** Predicted *Cp*MetRS amino acid sequences obtained prior to treatment and on experiment day 9. The calf number and day of experiment is indicated. The sequence from calf 74 day 1 (prior to treatment) is identical to the *Cp*MetRS in the cryptoDB genome database. The box highlights the predicted amino acid mutations in the *Cp*MetRS of parasites shed by calves 74 and 76 on day 9. **(b)** Sanger sequencing chromatograms and DNA sequences for oocysts shed by calves 74 and 76 on day 9.
