## Supplemental Figure 3 for "Spontaneous selection of *Cryptosporidium* drug resistance in a calf model of infection"

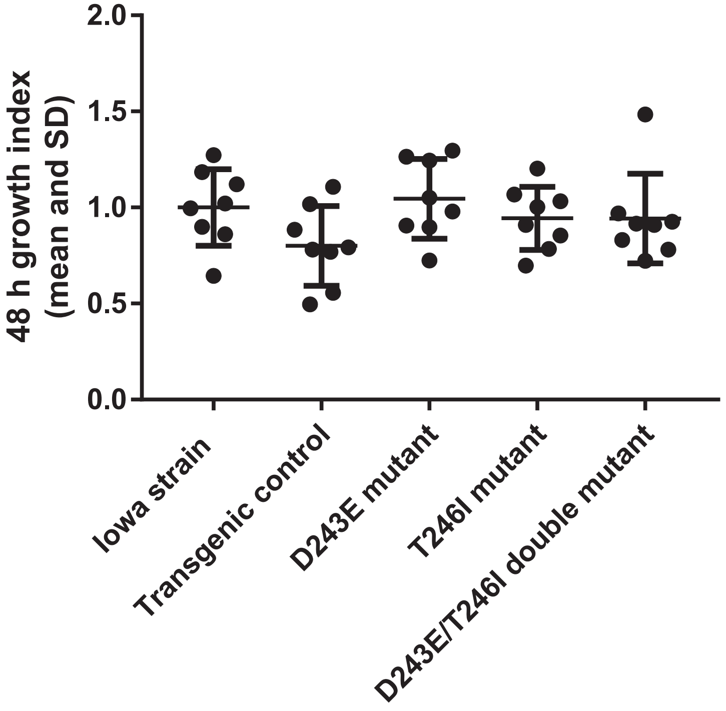


**Supplemental Fig. 3. In vitro growth of transgenic *C. parvum* lines relative to Iowa strain *C. parvum*.** HCT-8 cell monolayers were infected with each parasite line for 48 h, followed by enumeration of parasites and host cell nuclei by high content microscopy (*39*). Data are shown as a growth index, calculated as the percent of host cells infected normalized to the mean percent of host cells infected by Iowa strain parasites on the same assay plate (n=8 culture wells per *C. parvum* line; no significant differences by Kruskal-Wallis and Dunn's multiple comparisons tests).
