## Supplemental Table 1 for "Spontaneous selection of *Cryptosporidium* drug resistance in a calf model of infection"

**Supplemental Table 1. PCR primers.**

| | |
|---|---|
| ***C. parvum* 18s rRNA primers used for qPCR** | |
| **Cp18s forward** | 5’TAGAGATTGGAGGTTGTTCCT3’ |
| **Cp18s reverse** | 5’CTCCACCAACTAAGAACGGCC3’ |
| **Primers used for CRISPR/Cas9 generation of *CpMetRS* mutants** | |
| **Primers used to confirm genomic integration** | |
| **P1** | 5’TGAGGACTTATCTATAAGTAGAGC3’ |
| **P2** | 5’TGGATGTTAGGAAATTAACACTC3’ |
| **Primers used to amplify synthetic constructs** | |
| **Recoded WT forward** | 5’GGGGAGTTCCCGTTCCAAATGACCAGGAACATGTTATGTATGTTTGGTTTGACGCATTGACTAACTACATAACAGGCCTTAAGTGGG3’ |
| **D243E forward** | 5’GGGGAGTTCCCGTTCCAAATGACCAGGAACATGTTATGTATGTTTGGTTTGAAGCATTGACTAACTACATAACAGGCCTTAAGTGGG3’ |
| **T246I forward** | 5’GGGGAGTTCCCGTTCCAAATGACCAGGAACATGTTATGTATGTTTGGTTTGACGCATTGATTAACTACATAACAGGCCTTAAGTGGG3’ |
| **D243E / T246I forward** | 5’GGGGAGTTCCCGTTCCAAATGACCAGGAACATGTTATGTATGTTTGGTTTGAAGCATTGATTAACTACATAACAGGCCTTAAGTGGG3’ |
| **Synthetic construct reverse** | 5’AATTAAGATAAAAAGAAAAACTTAATCGATACTATCCTACACG3’ |
| **Additional primers used for cloning CRISPR constructs** | |
| **Eno-nluc-neo forward** | 5’GTAGGATAGTATCGATTAAGTTTTTCTTTTTATCTTAATTTGGGGAAACTAAATATACTGAAATTCGG3’ |
| **Eno-nluc-neo reverse** | 5’GTAAGCTATTTCAATTTATATTAAAGTTATATATATTTAACTAAATAACTTCAGAAGAATTCGTCAAGAAGACG3’ |
| **End primer forward** | 5’GGGGAGTTCCCGTTCCAAATG3’ |
| **End primer reverse** | 5’GTAAGCTATTTCAATTTATATTAAAGTTATATATATTTAACTAAATAACTTCAGAAG3’ |
| **Primers used for *CpMetRS* sequencing** | |
| **CpMetRS1-R** | 5’TTTGCCCATGTTCATCAGTTCC3’ |
| **CpMetRS2-R** | 5’ATAGATAAGTCCTCAGAATCGCTG3’ |
| **CpMetRS3-R** | 5’TATCATTCCCAAATCTCCCTTCTC3’ |
| **CpMetRS4-R** | 5’TCTATCCTCTCTTCATCAGACTTG3’ |
| **CpMetRS5-R** | 5’TTTCTCTGCTTTAAGCTCTTCG3’ |
| **CpMetRS6-R** | 5’CAGACTCAAGAAAGCTGGATG3’ |
| **Primers used to clone *CpMetRS* into AVA plasmid for recombinant expression** | |
| **CpMetRSAVAPCR-F** | 5’GGGTCCTGGTTCGATGACTAAGATAAATACAACATACAGAAACTCGTC3’ |
| **CpMetRSAVAPCR-R2** | 5’CTTGTTCGTGCTGTTTACTATTTTCCCTTATCTTTAGTTTCACTTAATTTCTTATTTCCT3’ |
